# GlioME: A Novel Organoid Model Retaining the Glioma Microenvironment for Personalized Drug Screening and Therapeutic Evaluation

**DOI:** 10.1101/2024.12.17.628908

**Authors:** Chengjun Zheng, Qiaodong Chen, Peng Wang, Delong Zhang, Zheng Fang, Yutong Feng, Jie Chen, Jiahong Chen, Yiwen Fu, Bao Yang, Qian Zhao, Fei Sun, Ying Zhang, Tao Jiang, Lemin Zheng, Zhaoshi Bao

## Abstract

Glioma is an aggressive brain tumor with a poor prognosis. Establishing an in vitro culture model that closely replicates the cellular composition and microenvironment of the original tumor has been challenging, limiting its clinical applications. Here, we present a novel approach to generate glioma organoids with a microenvironment (GlioME) from patient-derived glioma tissue. These organoids maintain the genetic and epigenetic characteristics of the primary tumor and preserve cell-to-cell interactions within the tumor microenvironment, including resident immune cells. Bulk RNA sequencing, whole exome sequencing, and DNA methylation analysis were used to confirm the molecular similarities between the organoids and primary glioma tissues. Immunofluorescence and flow cytometry were used to assess immune cell viability, comparing GlioME with floating glioma organoids. GlioME exhibited high responsiveness to chemotherapy and targeted therapy, demonstrating its potential for therapeutic screening applications. Notably, GlioME accurately predicted patient response to the recently approved MET inhibitor, vebreltinib. Thus, this organoid model provides a reliable in vitro platform for glioma microenvironment-related research and clinical drug screening.

## Introduction

Gliomas are the most common malignant brain tumors, characterized by their aggressive nature and poor prognosis^1^. Despite advancements in radiation and chemotherapy, the five-year survival rate for patients with glioma remains low^2^. A significant challenge is the lack of experimental models that accurately replicate the tumor microenvironment and cellular heterogeneity of human gliomas, limiting the understanding of disease mechanisms and the development of effective therapies.

Current experimental models include cell lines, patient-derived cell lines (PDCs), and patient-derived tumor xenografts (PDTX) ^3^. However, cell lines fail to represent the complexity of the original tumor, and PDCs are overly simplistic and lack the structural and physiological characteristics of tumor tissue. Moreover, studies have shown discrepancies between primary and original tumor cells. PDTX models, while useful, are time-consuming to establish, have low success rates, and cannot replicate the immune system. Patient-derived tumor organoids have recently emerged as promising three-dimensional in *vitro* systems^4^. These organoids can be established from small tissue samples in a short time, while retaining the histopathological and genomic characteristics of the original tumor^5^.

Several organoid models of glioblastoma have been developed. Researchers have used tumor fragments from patients and employed floating culture methods to create patient-derived glioblastoma organoids that can be passaged in *vitro*^6^. Others have used enzymatic digestion to break down tumor tissues into single cells, which are then cultured in Matrigel, allowing tumor cells to spontaneously form organoid structures^7^. Additionally, some researchers have used human induced pluripotent stem cells (iPSCs) combined with CRISPR-Cas9 technology to knock out genes, such as PTEN and TP53, generating glioblastoma-like organoids^8^. Despite these advancements, replicating the in vivo tumor microenvironment remains challenging^9^. Floating-culture organoids lose immune cells over time, and enzymatic digestion disrupts the complex cellular interactions within the tumor. Organoids derived from iPSCs lack non-tumor components. These limitations prevent these models from preserving the original tumor microenvironment, restricting their utility in studying glioblastoma. To date, no method has been developed that can fully retain the glioblastoma microenvironment in organoid models.

To address these limitations, we established glioma organoids that preserved the tumor’s immune microenvironment. Comprehensive histological, molecular, and genetic analyses confirmed that these organoids closely resemble the original tumors, maintaining key biomarkers, similar proportions of immune and stromal cells, and the heterogeneity of both tumor tissue and cells. We also compared this method with another approach using patient-derived tumor fragments that do not require single-cell dissociation to create glioma organoids. Furthermore, we explored drug screening applications, demonstrating that these organoids enable high-throughput drug testing, underscoring their potential significance in maintaining the tumor immune microenvironment for therapeutic research.

## Results

### Culture and characterization of patient-derived glioma organoids

To preserve cellular components and their interactions within the tumor, we developed a new glioma organoid culture protocol (GlioME) and successfully established an organoid culture system from patient samples of multiple glioma subtypes (Figure 1a). In this protocol, we directly inoculated mechanically minced fresh tumor tissue into the low-growth factor Matrigel and cultured it in a serum-free medium without additional growth factors, such as EGF and bFGF. We successfully established organoid cultures from the tumors of 30 out of 34 (88.2%) patients, covering three glioma subtypes and different WHO grades, including IDH-mutant samples and primary and recurrent tumors (Figure 1b). Additionally, we cultured floating glioma organoids (FGs) paired with each sample using mechanical tissue fragments without dissociation into single cells and performed a high-precision comparative analysis of the two methods.

**Figure 1:**
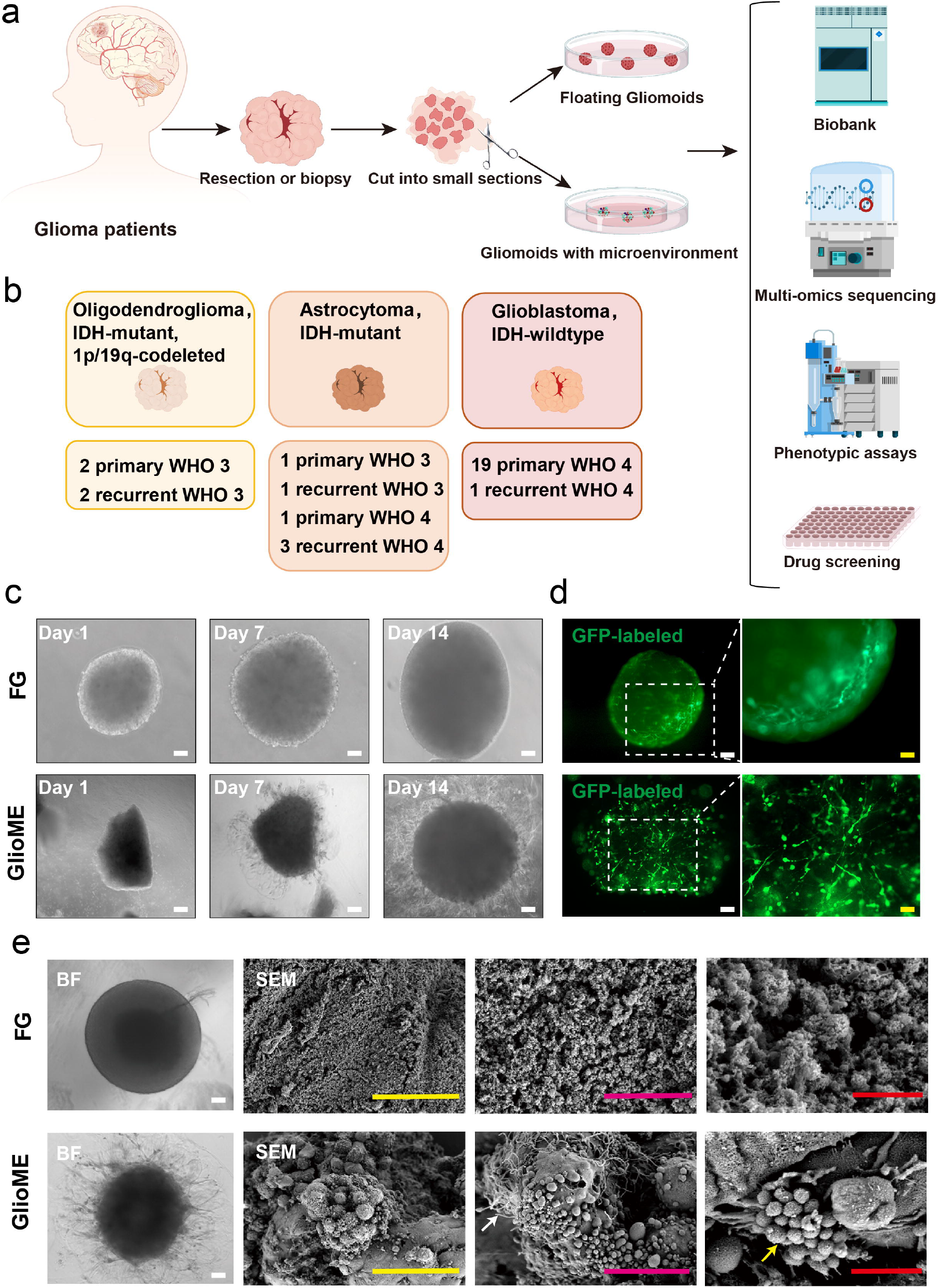
Establishment and morphological characterization of floating organoids and glioma organoids with microenvironment (GlioME). (**a**) Schematic representation of the cultivation strategy and characterization for floating and GlioME. (**b**) Overview of established glioma organoids across different subtypes and grades. (**c**) Representative bright-field microscopy images of floating and microenvironment-retaining glioma organoids at days 1, 7, and 14 of cultivation. (**d**) Fluorescent imaging of glioma organoids transduced with GFP-lentiviral particles. (**e**) Scanning electron microscopy (SEM) images of glioma organoids, with white arrows indicating tumor-associated macrophages and yellow arrows indicating exosomes.

Figure 1c shows the organoids observed under light microscopy, which formed a stable morphology after 1–2 weeks of culturing. Among the organoids produced by the two culture methods, the organoids with higher growth activity tended to be more spherical, while the FGs had a smoother surface due to culturing due to the use of an orbital shaker during cultivation. Figure 1c shows that GlioME cells extended along the matrix gel to form tentacle-like structures, reflecting the invasive growth characteristics of glioma cells in vivo. Figure 1d shows that after transfection with a lentivirus carrying GFP, the interaction between cells within the organoid was more complex than that seen in two-dimensional culture. Figure 1e provides a detailed image through SEM, showing that GlioME organoids have stronger heterogeneity in their components, retain tumor-associated macrophages, and display exosomes secreted by the cells.

### Histological and immunohistochemical comparison of glioma organoids and original tumors

All established organoids successfully reproduced the histopathological characteristics and molecular patterns of the original tumors. Figure 2a shows hematoxylin-eosin (HE) staining of three different glioma subtypes cultured using the two methods after two weeks of culture (Figure 2a). The HE staining results showed that FGs and GlioMEs were morphologically similar in glioblastomas, but there were certain differences in IDH-mutant gliomas. For example, in an IDH-mutant astrocytoma, the GlioMEs better retained the characteristics of gemistocytes, whereas the FGs were relatively large and showed a large number of proliferating granular cells. This discrepancy could be due to the slower proliferation of IDH-mutant glioma cells. The floating method tends to promote excessive proliferation of specific cell types, resulting in inconsistencies in the cellular composition compared to the original tumor. In glioblastomas, organoids from both culture methods retained the characteristics of abnormal nuclear division, heterogeneous tumor cells, and a dense cellular arrangement, resulting in a chaotic distribution pattern.

**Figure 2:**
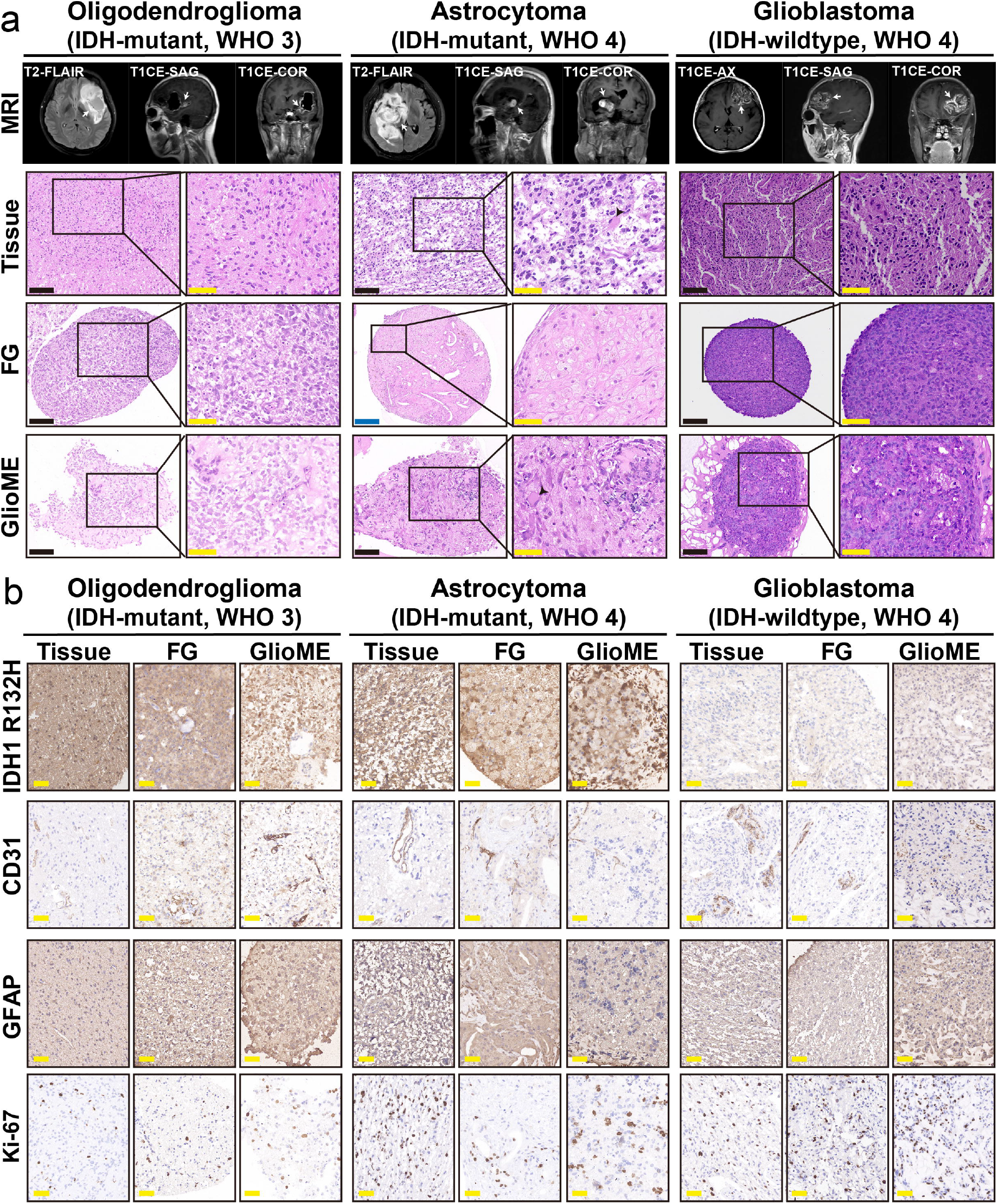
Preservation of heterogeneous histology and tumor microenvironment features in glioma organoids. (**a**) Contrast-enhanced T1 MRI showing sampling site (indicated by white arrow) and corresponding tumor tissue with hematoxylin and eosin (HE) staining of patient-derived glioma organoids after two weeks of cultivation. (**b**) Concordant expression of IDH1 R132H, CD31, GFAP, and KI-67 in astrocytoma, oligodendroglioma, glioblastoma, and their derived organoids.

Subsequently, key tumor markers in the original tumor tissue and glioma organoids were compared using HE staining and specific antibody labeling (Figure 2b). FGs and GlioMEs exhibited similar staining patterns to the paired parental tumors for markers, such as IDH and GFAP. Tumor-infiltrating vascular endothelial cells labeled with CD31 were retained in both organoid types. KI-67 staining showed that both culture methods retained a considerable proportion of actively proliferating cells within two weeks, maintaining a proliferation rate similar to that of the parental tumor.

### Genomic profiling and molecular consistency of glioma organoids

To verify whether the organoids retained the genetic characteristics of the parental tumor, we compared the genomic consistency between the organoids and the parental tumor using bulk RNA sequencing, whole exome sequencing, and DNA methylation analysis. Due to sample size limitations, not all organoids and parental tumors were sequenced. Finally, we performed bulk RNA sequencing on samples from six patients, whole-exome sequencing on samples from four patients, and DNA methylation sequencing on samples from five patients.

Although there were differences between the organoids from different patients, the RNA-seq results from the two types of organoids and the original tumor tissues showed a high correlation (Figure 3a). Subsequently, we performed unsupervised clustering analysis on 18 samples from six patients using 1,421 tumor-related genes^11^, and the results showed that tissues and organoids from the same patient had specific clustering (Figure 3b). Whole-exome sequencing of GlioMEs and parental tumor samples revealed CNVs and single nucleotide mutations commonly found in gliomas (Figure 3c-d). As expected, GlioMEs successfully retained the mutant genes of the parental tumor, with most CNVs observed in the parental tumors being retained in the organoids (Figure 3c). The mutation spectra of GlioMEs and parental tumors showed approximately 76.7% similarity (Figure 3d).

**Figure 3:**
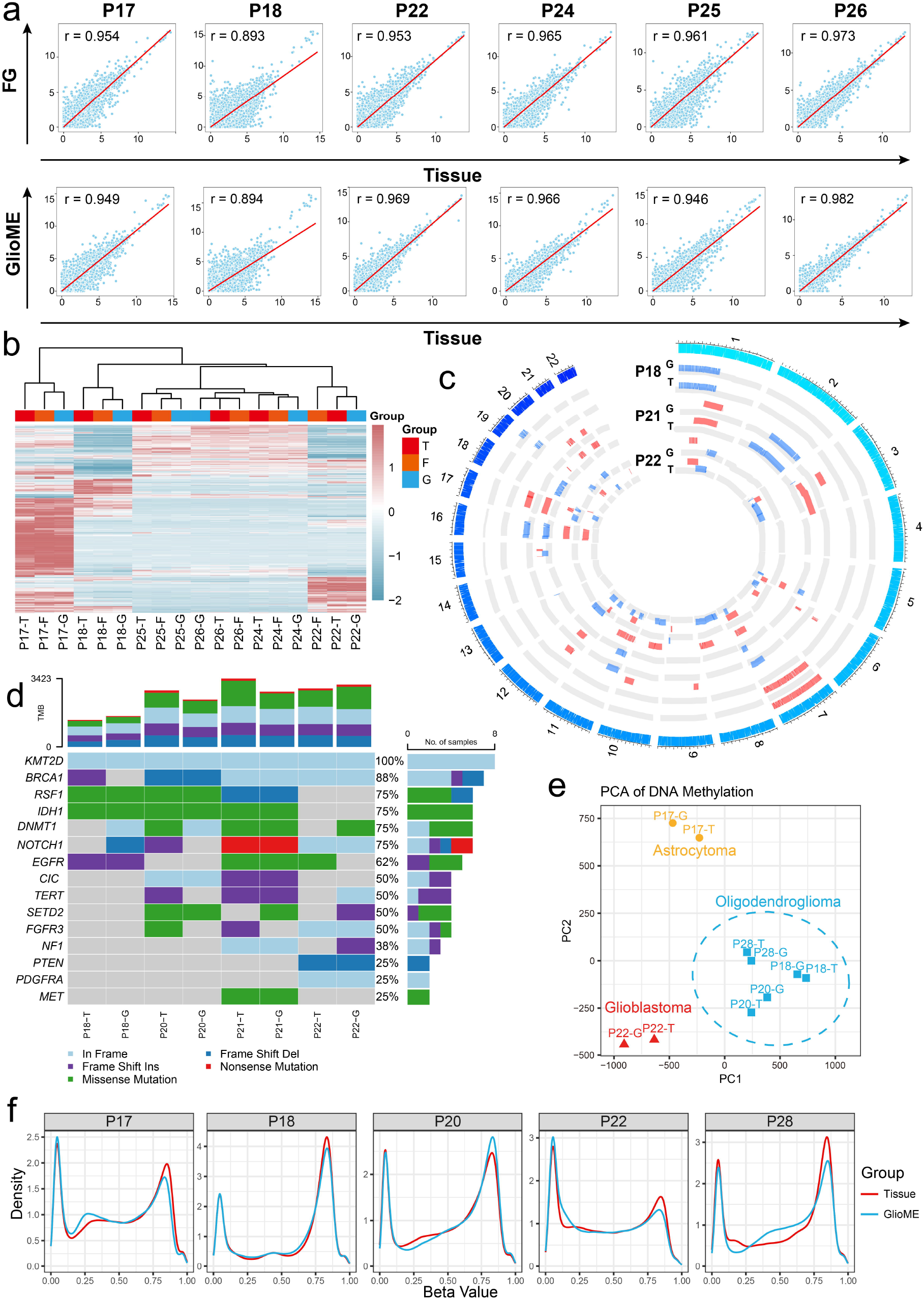
Glioma organoids maintain consistency with the original tumor tissues at the transcriptome, methylation, and exon mutation spectrum levels. (**a**) Pearson correlation plot of bulk-RNAseq gene expression data, comparing the gene expression correlation between the original tissue and the two types of glioma organoids (FG and GlioME). r: correlation coefficient. (**b**) Unsupervised clustering of 1,421 brain cancer-related genes based on bulk RNA sequencing data of 18 samples from six patients, showing the consistency of the transcriptome between the two glioma organoids and their corresponding original tumor tissues. (**c**) Circle plot of the CNV spectrum obtained by whole exome sequencing, showing the similarity in CNV characteristics between GlioMEs (G) and their corresponding original tumor tissues (T). (**d**) Mutation spectrum of commonly mutated genes, showing the somatic mutation characteristics in GlioMEs and their original tumor tissues. (**e**) Principal component analysis clustering plot, showing that GlioME organoids and their corresponding patient-derived tumor tissues are clustered together, while showing subtype specificity. (**f**) Density plot of DNA methylation beta values, comparing tumor tissues and GlioMEs derived from five patients.

In addition, we studied the DNA methylation levels in organoids and tumor tissues. Principal component analysis showed that the tumor tissues and GlioMEs from the three oligodendrogliomas clustered together, with a higher similarity observed within the same subtype compared to different subtypes (Figure 3e). DNA methylation patterns in tumors and GlioMEs from the same patient showed a high degree of similarity (Figure 3f). In summary, these results show that GlioMEs cultured in *vitro* effectively retain the genomic, transcriptomic, and DNA methylation patterns of the original tumors.

### Immune microenvironment preservation in glioma organoids

To further investigate the heterogeneity of cells within the organoids, we analyzed bulk RNA sequencing data and used the Cibersort-ABS algorithm to deconvolute immune cell components in the parental tumor and organoids^12^. We first focused on tumor-associated macrophages, which comprise the largest proportion in gliomas, and found that, after two weeks of culture, GlioMEs better preserved the cellular composition of the original tumor tissues (Figure 4a).

**Figure 4:**
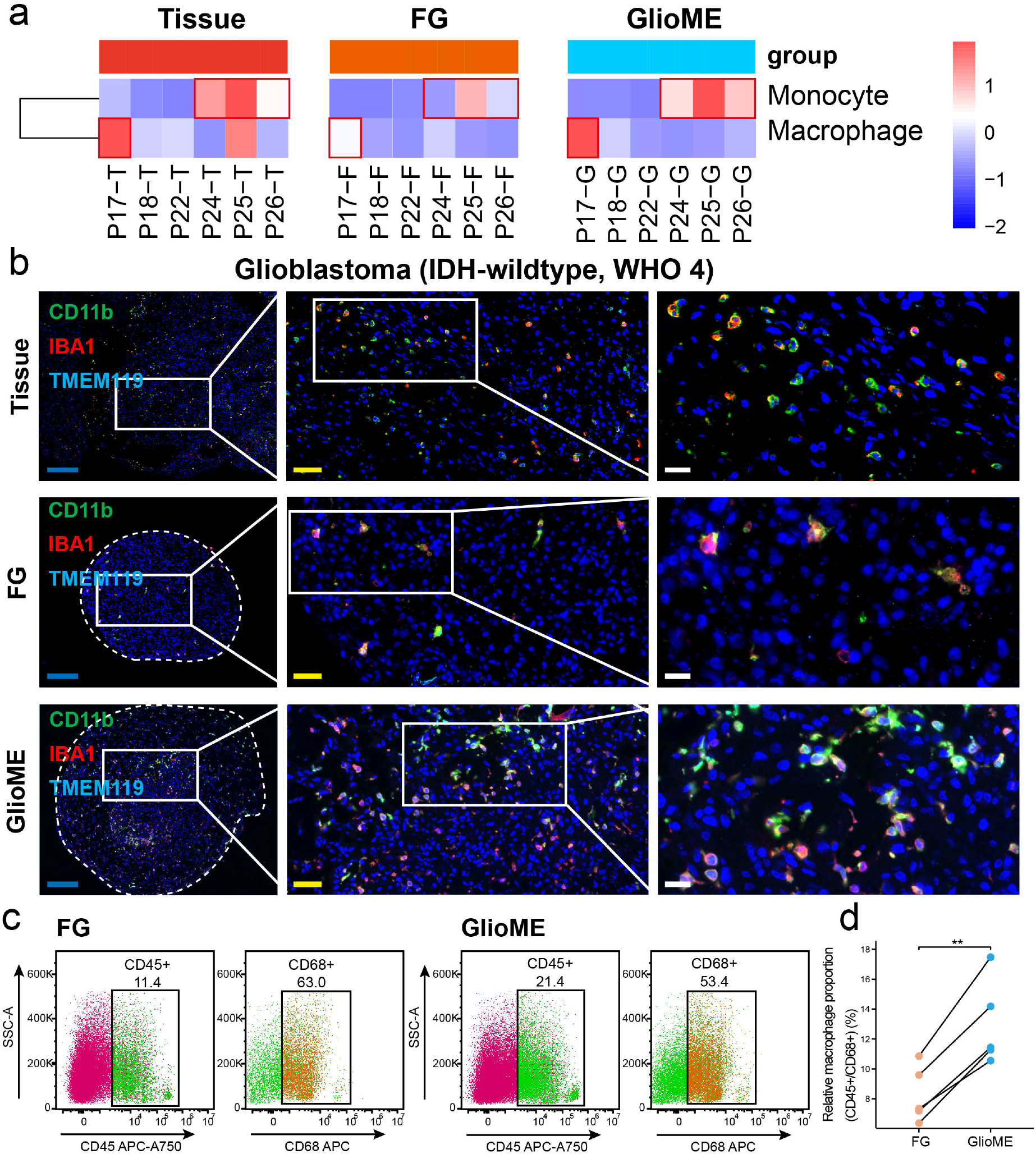
Enhanced retention of macrophages in GlioMEs compared to FGs. (**a**) Proportions of macrophages and monocytes in each sample estimated by CIBERSORT-ABS deconvolution analysis of bulk RNA sequencing data. (**b**) Immunofluorescence analysis showing the expression of CD11b, IBA1, and TMEM119 markers in tumor tissue, and in FGs and GlioMEs after two weeks of cultivation. (**c**) FACS analysis of CD45+ and CD68+ cells in FGs and GlioMEs after two weeks of cultivation. (**d**) FACS quantification of CD45+ and CD68+ cells in FGs and GlioMEs after two weeks of cultivation.

We then labeled CD11b+, IBA1+, and TMEM119+ macrophages and microglia by immunofluorescence staining after two weeks of culture. We compared the tumor-associated macrophages in the parental tumor and the two types of organoids from the same patient. The results showed that the number of macrophages retained in the GlioMEs was significantly higher than that in the FGs (Figure 4b). Subsequently, we performed flow cytometry on the two organoids cultured for two weeks in 10 patients (Figure 4d) and found that the number of CD45+/CD68+ macrophages in GlioMEs was approximately twice that in FGs (Figure 4f).

Although T cell infiltration is relatively rare in glioma tissue, we compared the two organoids with T cells from the original tumor. Using the ssGSEA algorithm to assess the expression levels of T cell-related genes, we found that the gene expression of GlioMEs more closely matched that of the parental tumors (Figure 5a). Analysis of gene expression levels for T-cell markers IFNG, FOXP3, GZMA, and GZMH confirmed this observation, showing that GlioMEs closely mirror the immune characteristics of the parental tumor based on bulk RNA sequencing (Figure 5b). Immunofluorescence staining showed that GlioMEs retained more CD4+ and CD8+ T cells than FGs (Figure 5c). Flow cytometry results showed that the average number of CD45+/CD3+ cells was higher in GlioMEs than in FGs, indicating that T cells were more difficult to retain in FGs (Figure 5d-e). Unlike other methods that involve the exogenous addition of immune cells, GlioMEs retain the original immune cells from the tumor tissue instead of reconstructing them. This allows for the effective preservation of the tumor’s immune microenvironment characteristics.

**Figure 5:**
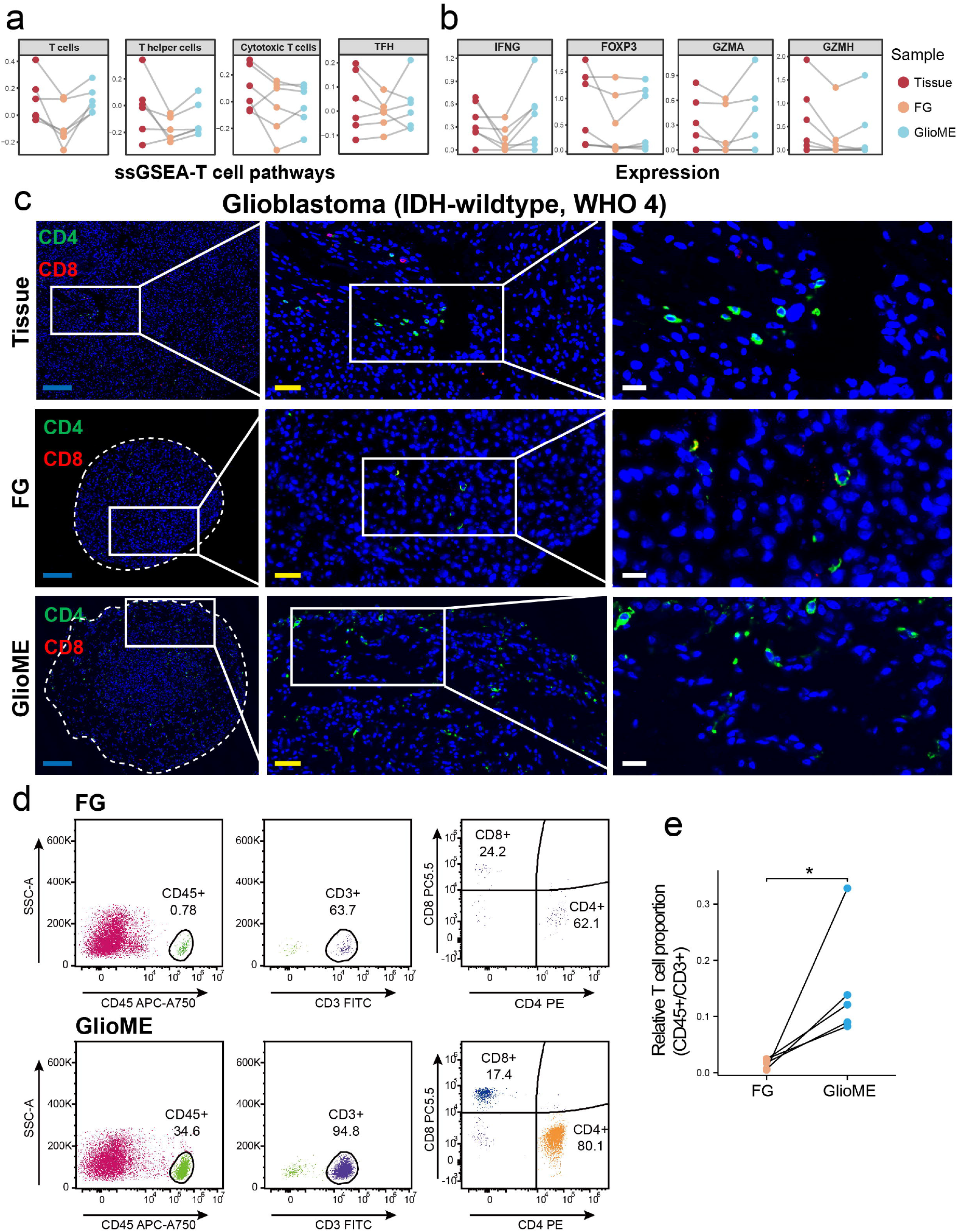
Enhanced retention of T cells in GlioMEs compared to FGs. (**a**) ssGSEA scores for four T cell-associated pathways based on bulk RNA sequencing in tissues, FGs, and GlioMEs. (**b**) Gene expression levels of IFNG, FOXP3, GZMA, and GZMH in tissues, FGs, and GlioMEs based on bulk RNA sequencing. (**c**) Immunofluorescence analysis showing the expression of CD4 and CD8 markers in tumor tissue, and in FGs and GlioMEs after two weeks of cultivation. (**d**) FACS analysis of CD45+/ CD3+/CD4+/CD8+ cells in FGs and GlioMEs after two weeks of cultivation. (**e**) FACS quantification of CD45+ and CD3+ cells in FGs and GlioMEs after two weeks of cultivation.

### Drug screening and clinical implication by GlioME

To evaluate the response of GlioMEs to immunotherapy, chemotherapy, and targeted therapy, we selected pexidartinib (a CSF1R inhibitor targeting M2 macrophages), temozolomide (the primary chemotherapy used in glioma treatment), and vebreltinib (a MET inhibitor). Due to its limited mechanism of action, the application of pexidartinib in glioma treatment is currently limited to animal models^13^. We performed immunofluorescence staining on organoids cultured for two weeks and treated for 6 days with 1 μM of pexidartinib. The results showed a significant reduction in the number of macrophages (Figure 6a), and flow cytometry further confirmed that the proportion of CD206+ macrophages was reduced (Figure 6b).

**Figure 6:**
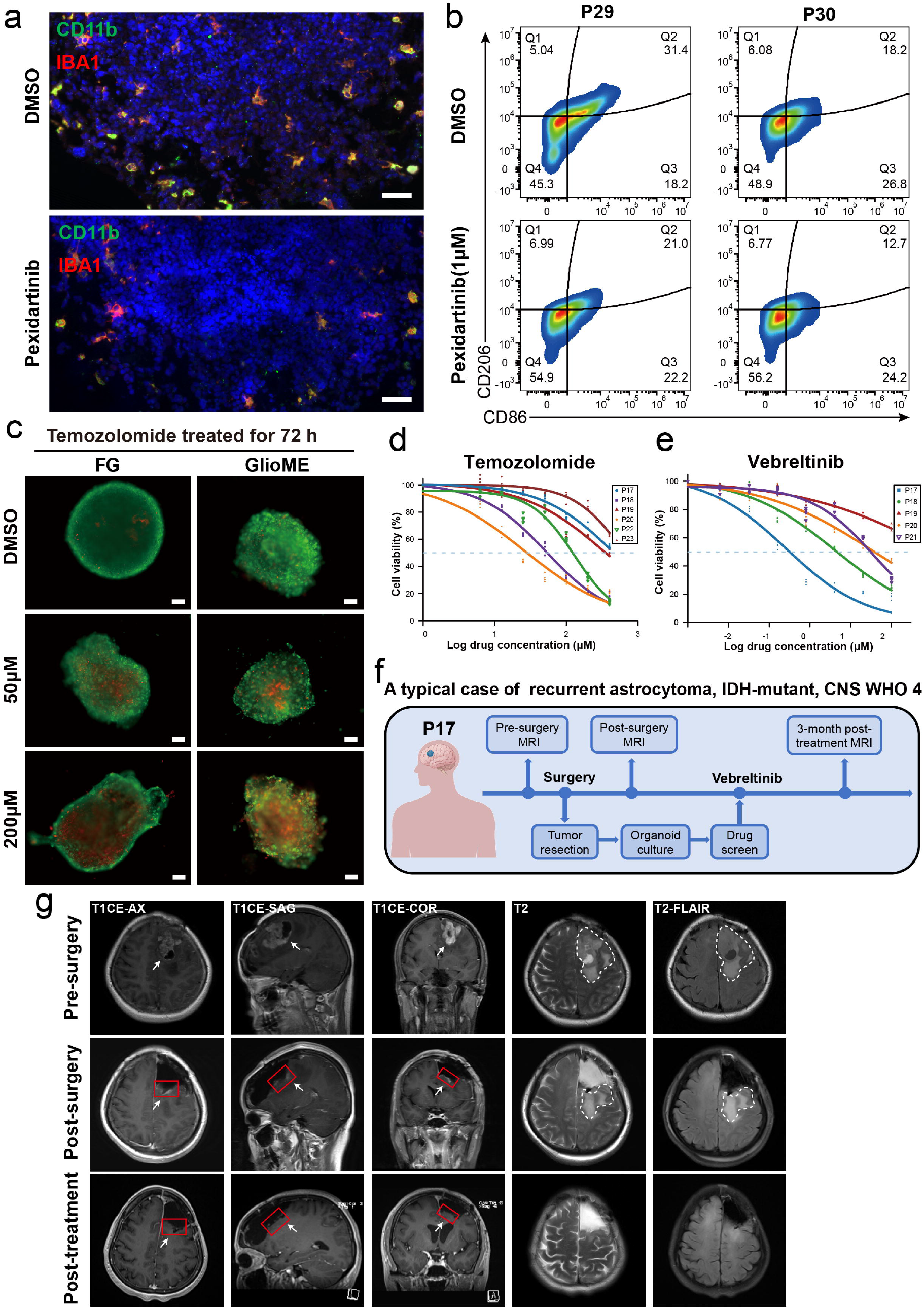
GlioME as a robust preclinical model for personalized drug screening. (**a**) Immunofluorescence staining of CD11b and IBA1 in GlioMEs treated with Pexidartinib (1 μM) for 6 days, with DMSO-treated organoids as controls. (**b**) Flow cytometry analysis of GlioME organoids treated with Pexidartinib (1 μM) for 6 days, using the gating strategy of CD45+/CD68+ to assess CD86 and CD206 expression; DMSO-treated organoids served as controls. (**c**) Dose-response curves of GlioMEs after 72 hours of temozolomide treatment. (d) Dose-response curves of GlioMEs after 72 hours of vebreltinib treatment. (**e**) Fluorescence images of GlioMEs stained with AO/PI after 72 hours of treatment with DMSO, 50 μM, and 200 μM temozolomide. (**f**) Flowchart of procedures and treatments for patient P17 with MET amplification. (**g**) MRI images of patient P17 before surgery, after surgery, and 2 months after vebreltinib treatment (3 months post-surgery). White arrows indicate the residual enhancing area post-surgery.

Temozolomide remains the standard chemotherapeutic regimen for glioma treatment^14^. Both organoid models reflected the patient’s sensitivity to temozolomide. We performed drug sensitivity testing on organoids derived from the same patient and used AO/PI staining to assess cell viability and death. The results showed a clear dose-response gradient, indicating that temozolomide effectively penetrated the organoids and exerted its therapeutic effects (Figure 6c). We also conducted drug sensitivity screening on GlioMEs from six patients, and the fitted dose-response curve showed differences in sensitivity to temozolomide and vebreltinib among different patients (Figure 6d and e).

We found that a patient with recurrent left frontal IDH-mutant astrocytoma, CNS WHO 4, showed greater sensitivity to vebreltinib than to temozolomide (Figure 6e). The patient was subsequently treated with vebreltinib for two months (Figure 6f). MRI images taken 3 months after treatment showed that vebreltinib achieved a significant clinical response, with the residual enhancing lesions disappearing and a reduction in the T2 flair lesion, along with symptom relief, resulting in complete remission (Figure 6g). Fluorescence in situ hybridization analysis), revealed MET amplification in the patient, which likely explains their sensitivity to vebreltinib.

## Discussion

Gliomas pose a significant challenge in oncology due to their significant heterogeneity and complex microenvironment, resulting in a lack of a universal treatment plan that can effectively target all gliomas^15,16^. The glioma microenvironment is composed of multiple cellular components, including tumor, immune, and stromal cells^17^. This complex cellular environment leads to the high heterogeneity of gliomas and drug resistance issues, making it difficult for treatments to yield consistent outcomes^18,19^. Patient heterogeneity in terms of genetic background, tumorigenesis mechanisms, and tumor molecular characteristics leads to significant differences in the efficacy of the same treatment regimen^20^. Additionally, within a single patient, gliomas show significant heterogeneity. Cells in different tumor areas have different phenotypes and genetic mutations, which means that some tumor cells may be sensitive to drugs, whereas others are resistant^21^. Our novel organoid culture method successfully cultured IDH-mutated grade 4 astrocytomas and organoids of other glioma subtypes, providing a reliable in vitro experimental model for glioma heterogeneity research. Tumor-infiltrating vascular endothelial cells labeled with CD31 were retained in both organoids. However, due to the lack of blood circulation, similar to that in vivo, the vascular cavity collapsed, leaving only endothelial cells. This remains a challenge in organoid research that must be addressed in the future.

The glioma microenvironment contains a large number of immune cells, such as tumor-associated macrophages and T cells. Tumors can recruit and reshape immune cells, transforming them into cells that support tumor growth and inhibit antitumor immune responses^22^, thereby more accurately reproducing the tumor microenvironment in vitro^23,24^. Currently, most methods for constructing organoids from patient tumor tissues use the enzymatic dissociation of tumor tissues into single cells^7^. However, organoids established using these methods fail to retain the original composition of the parental tissue or the tumor microenvironment. Retaining tumor cells, autologous tumor-related immune cells, and stromal cells are crucial for drug screening and reflecting the in vivo situation^20^. To address these challenges, we proposed a new organoid culture method for primary and recurrent gliomas and successfully constructed organoid models of primary and recurrent gliomas. This protocol does not require tissue dissociation into single cells and can preserve cell-to-cell connections and non-tumor components, such as immune cells and endothelial cells.

As the culture time progresses, the morphological differences between successfully cultured organoids and necrotic tissues gradually become more pronounced. Necrotic tissues do not grow but gradually shrink and lose their original morphology, whereas organoids grow along the matrix gel to form a three-dimensional structure. The success rate of organoid culture is mainly affected by the activity of the tumor tissue, sampling site, and transportation time, with tumor activity and sampling site being particularly critical. The tumor density in the MRI-enhanced area is high, and the survival rate is high; for glioblastoma, sampling from the necrotic core area should be avoided. The surgeon must carefully select the area with the densest tumor under the microscope, and the sampling process depends on close cooperation between the collector and surgeon. During the culture process, we avoided the use of exogenous EGF/bFGF and serum but chose a matrix gel with a low growth factor content to reduce the effect of growth factors on the proliferation ability of specific cells, thereby retaining a cell ratio similar to that of the original tumor. The results of HE staining, immunohistochemistry, whole exome sequencing, DNA methylation, and transcriptome sequencing showed that the cultured organoids were highly similar to the parent tumor at both histological and molecular levels. Unlike organoid cultures obtained by floating methods, our method specifically retains immune cells infiltrating the tumor tissue, including macrophages and T cells, rather than co-culturing with exogenously added immortalized immune cells. Immunofluorescence, flow cytometry, and gene sequencing revealed that the organoids retained an immune microenvironment consistent with that of the original tumor.

Our findings highlight the unique advantages of GlioMEs as innovative organoid models. Unlike traditional cell lines and animal models, GlioMEs retain the immune microenvironment, allowing faster and more accurate immunotherapy screening. For instance, treatment with the CSF1R inhibitor pexidartinib led to a clear and rapid reduction in the number of macrophages^13^. This provides a straightforward and efficient method to assess the drug’s effects.

Furthermore, the complex layers of glioma tissue make treatment challenging, increasing the possibility of drug resistance and recurrence. To address this critical clinical issue, a sufficiently accurate personalized treatment platform that can effectively predict and respond to each patient’s drug-resistance risk and treatment response is urgently required. In this study, we performed drug screening based on a newly invented organoid system to test the effects of temozolomide and vebreltinib and then applied the screening results to the clinical treatment of a patient. The accurate prediction of vebreltinib sensitivity in a patient with MET amplification demonstrates the reliability of GlioMEs for drug screening. Although MET amplification is not currently an approved indication for vebreltinib^25,26^, our GlioME approach successfully identified a patient who could benefit from this drug. This suggests that GlioMEs not only provide drug guidance for glioma patients but also serve as an effective platform for verifying the potential expansion of drug indications.

In summary, our results demonstrate that GlioME can retain the parental glioma microenvironment in vitro and reliably test various therapeutic drugs. This study validates the ability of GlioME to maintain stability in culture for at least one month, which provides a solid foundation for further experiments. This method not only provides an important tool for basic research on gliomas but also helps in the formulation of personalized treatment plans. In the future, we will continue to optimize and expand this technology to promote advances in glioma-related research and clinical treatments. This organoid model will be used to further characterize the dynamic changes in the tumor microenvironment, explore the role of immune cells, such as T cells and macrophages, in tumor killing, and assess the response of these cells to different treatment regimens.

## References

1. Ostrom, Q. T. et al. CBTRUS Statistical Report: Primary Brain and Other Central Nervous System Tumors Diagnosed in the United States in 2011–2015. Neuro-Oncol. 20, iv1–iv86 (2018).

2. Stupp, R. et al. Radiotherapy plus Concomitant and Adjuvant Temozolomide for Glioblastoma. N. Engl. J. Med. 352, 987–996 (2005).

3. Robertson, F. L., Marqués-Torrejón, M.-A., Morrison, G. M. & Pollard, S. M. Experimental models and tools to tackle glioblastoma. Dis. Model. Mech. 12, dmm040386 (2019).

4. Wu, W., Li, X. & Yu, S. Patient-derived Tumour Organoids: A Bridge between Cancer Biology and Personalised Therapy. Acta Biomater. 146, 23–36 (2022).

5. Drost, J. & Clevers, H. Organoids in cancer research. Nat. Rev. Cancer 18, 407–418 (2018).

6. Jacob, F. et al. A Patient-Derived Glioblastoma Organoid Model and Biobank Recapitulates Inter- and Intra-tumoral Heterogeneity. Cell 180, 188–204.e22 (2020).

7. Hubert, C. G. et al. A Three-Dimensional Organoid Culture System Derived from Human Glioblastomas Recapitulates the Hypoxic Gradients and Cancer Stem Cell Heterogeneity of Tumors Found In Vivo. Cancer Res. 76, 2465–2477 (2016).

8. Wang, C. et al. A multidimensional atlas of human glioblastoma-like organoids reveals highly coordinated molecular networks and effective drugs. Npj Precis. Oncol. 8, 19 (2024).

9. Klemm, F. & Joyce, J. A. Microenvironmental regulation of therapeutic response in cancer. Trends Cell Biol. 25, 198–213 (2015).

10. Louis, D. N. et al. The 2021 WHO Classification of Tumors of the Central Nervous System: a summary. Neuro-Oncol. 23, 1231–1251 (2021).

11. Zhao, M., Liu, Y., Ding, G., Qu, D. & Qu, H. Online database for brain cancer-implicated genes: exploring the subtype-specific mechanisms of brain cancer. BMC Genomics 22, 458 (2021).

12. Newman, A. M. et al. Determining cell type abundance and expression from bulk tissues with digital cytometry. Nat. Biotechnol. 37, 773–782 (2019).

13. Pyonteck, S. M. et al. CSF-1R inhibition alters macrophage polarization and blocks glioma progression. Nat. Med. 19, 1264–1272 (2013).

14. Tan, A. C. et al. Management of glioblastoma: State of the art and future directions. CA. Cancer J. Clin. 70, 299–312 (2020).

15. van den Bent, M. J. et al. Primary brain tumours in adults. Lancet Lond. Engl. 402, 1564–1579 (2023).

16. Osuka, S. & Van Meir, E. G. Overcoming therapeutic resistance in glioblastoma: the way forward. J. Clin. Invest. 127, 415–426 (2017).

17. Vitale, I., Manic, G., Coussens, L. M., Kroemer, G. & Galluzzi, L. Macrophages and Metabolism in the Tumor Microenvironment. Cell Metab. 30, 36–50 (2019).

18. Nicholson, J. G. & Fine, H. A. Diffuse Glioma Heterogeneity and Its Therapeutic Implications. Cancer Discov. 11, 575–590 (2021).

19. Jayaram, M. A. & Phillips, J. J. Role of the Microenvironment in Glioma Pathogenesis. Annu. Rev. Pathol. 19, 181–201 (2024).

20. Goenka, A. et al. The Many Facets of Therapy Resistance and Tumor Recurrence in Glioblastoma. Cells 10, 484 (2021).

21. Tomar, M. S., Kumar, A., Srivastava, C. & Shrivastava, A. Elucidating the mechanisms of Temozolomide resistance in gliomas and the strategies to overcome the resistance. Biochim. Biophys. Acta Rev. Cancer 1876, 188616 (2021).

22. Joseph, J. V., Blaavand, M. S., Daubon, T., Kruyt, F. A. & Thomsen, M. K. Three-dimensional culture models to study glioblastoma - current trends and future perspectives. Curr. Opin. Pharmacol. 61, 91–97 (2021).

23. Gangoso, E. et al. Glioblastomas acquire myeloid-affiliated transcriptional programs via epigenetic immunoediting to elicit immune evasion. Cell 184, 2454–2470.e26 (2021).

24. Liu, I., Hack, O. A. & Filbin, M. G. The imitation game: How glioblastoma outmaneuvers immune attack. Cell 184, 2278–2281 (2021).

25. Hu, H. et al. Mutational Landscape of Secondary Glioblastoma Guides MET-Targeted Trial in Brain Tumor. Cell 175, 1665–1678.e18 (2018).

26. Bao, Z. et al. PTPRZ1-METFUsion GENe (ZM-FUGEN) trial: study protocol for a multicentric, randomized, open-label phase II/III trial. Chin. Neurosurg. J. 9, 21 (2023).

